# Speed, accuracy, sensitivity and quality control choices for detecting clinically relevant microbes in whole blood from patients

**DOI:** 10.1101/549477

**Authors:** James Thornton, George S. Watts, Ken Youens-Clark, Lee D. Cranmer, Bonnie L. Hurwitz

## Abstract

Infections are a serious health concern worldwide, particularly in vulnerable populations such as the immunocompromised, elderly, and young. Advances in metagenomic sequencing availability, speed, and decreased cost offer the opportunity to supplement or replace culture-based identification of pathogens with DNA sequence-based diagnostics. Adopting metagenomic analysis for clinical use requires that all aspects of the pipeline are optimized and tested, including data analysis. We tested the accuracy, sensitivity, and resource requirements of Centrifuge within the context of clinically relevant bacteria. Binary mixtures of bacteria showed Centrifuge reliably identified organisms down to 0.1% relative abundance. A staggered mock bacterial community showed Centrifuge outperformed CLARK while requiring less computing resources. Shotgun metagenomes obtained from whole blood in three febrile neutropenia patients showed Centrifuge could identify both bacteria and viruses as part of a culture-free workflow. Finally, Centrifuge results changed minimally by eliminating time-consuming read quality control and host screening steps.

**AUTHOR SUMMARY:** Immunocompromised patients, such as those with febrile neutropenia (FN), are susceptible to infections, yet cultures fail to identify causative organisms ~80% of the time. High-throughput metagenomic sequencing offers a promising approach for identifying pathogens in clinical samples. Mining through metagenomes can be difficult given the volume of reads, overwhelming human contamination, and lack of well-defined bioinformatics methods. The goal of our study was to assess Centrifuge, a leading tool for the identification and quantitation of microbes, and provide a streamlined bioinformatics workflow real-word data from FN patient blood samples. To ensure the accuracy of the workflow we carefully examined each step using known bacterial mixtures that varied by genetic distance and abundance. We show that Centrifuge reliably identifies microbes present at just 1% relative abundance and requires substantially less computer time and resource than CLARK. Moreover, we found that Centrifuge results changed minimally by quality control and host-screening allowing for further reduction in compute time. Next, we leveraged Centrifuge to identify viruses and bacteria in blood draws for three FN patients, and confirmed suspected pathogens using genome coverage plots. We developed a web-based tool in iMicrobe and detailed protocols to promote re-use.

## INTRODUCTION

The current gold standard for clinical diagnosis of infections relies on isolating organisms by culture-based methods followed by identification and drug resistance testing. Methods for identifying pathogens that rely on culture have several drawbacks including fastidious bacteria, the time required for growth in culture, and the difficulty targeting viruses, fungi, and parasites. Identifying pathogens directly from biological samples by DNA sequencing can overcome the above limitations of culture and may improve the rate and speed of diagnosis. For these reasons, metagenomic shotgun sequencing of pathogens has been referred to as the holy grail of infection diagnosis (Ecker et al., 2010). While culturing samples is the current standard for infection diagnosis, it can have a high failure rate in some scenarios. For example, a study examined the problem of culture-based diagnosis of infection in febrile neutropenia and found that only ~16% (609 of 3,756) febrile neutropenia patients were culture positive (van Walraven & Wong, 2014). Also, the hazard ratio of dying was nearly four-fold higher in culture-negative patients than for patients where no culture was taken (presumably due to lack of fever), indicating the high cost in lives when cultures fail. Therefore, we seek to apply metagenomic sequencing to overcome the low rate and time delay of culture-based diagnostic methods in clinical settings such as febrile neutropenia.

The potential of metagenomic shotgun sequencing has been demonstrated in a broad range of infection scenarios including: leptospirosis (Wilson et al., 2014), nosocomial transmission of a drug-resistant bacteria (Snitkin et al., 2012), foodborne illness (Ashton et al., 2015), and infectious disease outbreaks (Quick et al., 2016). Despite successes using metagenomic shotgun sequencing to identify pathogens, routine application in clinical settings will require accurate, efficient classification, with minimized sample contamination. For example, while a small group of studies have reported on high-throughput metagenomic sequencing for identifying pathogens from immunocompromised patients where samples were not enriched for microbes, resulting in less than 1% of reads being pathogen-specific (Naccache et al., 2014; Parize et al., 2017) and dramatically reducing the diagnostic possibilities from the data (Frey et al., 2014). To begin addressing these inefficiencies, we developed an approach to increase the proportion of pathogen-derived reads in samples and applied it to the patient samples reported here.

On the data analysis side, there are no standards for analysis of metagenomic data obtained from clinical samples; however, there have been recent innovations in taxonomic classification algorithms that make it possible to quantify microbial species directly from reads in metagenomic datasets rapidly. These algorithms use two main approaches to assign reads to species in a reference database including: (1) a mapping approach using a Burrows-Wheeler transform (Li & Durbin, 2009; M. Burrows, 1994) used by Centrifuge (Kim, Song, Breitwieser, & Salzberg, 2016) or (2) a pseudo-alignment approach based on discriminating k-mers used by CLARK (Ounit, Wanamaker, Close, & Lonardi, 2015a). These algorithms outperform local alignment methods concerning both speed and capacity and can, therefore, better handle the number of reads in metagenomes (Bazinet & Cummings, 2012; Ounit, Wanamaker, Close, & Lonardi, 2015b; Rosen, Reichenberger, & Rosenfeld, 2011; Wood & Salzberg, 2014). However, comparisons between these algorithmic approaches to determine the accuracy of taxonomic assignment in clinically relevant metagenomes are lacking.

Here we report the accuracy and sensitivity of Centrifuge utilizing defined clinically relevant samples, compare its performance to CLARK, and finally analyze datasets obtained from patients following depletion of human cells to enrich for pathogen DNA. Lastly, we test the effect of excluding quality control and host-screening by alignment on the classification of reads by Centrifuge. This work provides a foundation for analysis of metagenomic data from clinical samples enriched for pathogens which use open-source software, requires a minimal computational resource, and provides rapid and accurate identification of pathogens. Our approach is freely available as web-based Apps in iMicrobe. Further, we provide the source code in GitHub: https://github.com/hurwitzlab/Centrifuge_HPC under the GNU open source license.

## RESULTS

### Centrifuge accuracy and sensitivity in controlled mixtures of bacteria

Because closely related clinically important bacteria can have diametric clinical consequences, (e.g., *E. coli* is a normal commensal while *S. flexneri* causes dysentery), we sought to test Centrifuge’s appropriateness as a tool for analyzing clinically relevant bacterial sequence datasets. We tested the linearity and threshold for detection of Centrifuge using three sets of bacterial mixtures, selected to represent taxonomic distances from phylum to genus-level. We created dilution mixtures over a six-log range of relative abundance with each organism ranging from 0.1% to 99.9% of the mixture (Figure 1). Centrifuge correctly identified all four species in the mixtures and misidentified less than one percent of the reads in any of the 18 combinations sequenced (false positives, Figure 1). Centrifuge was sensitive to the lowest relative abundance (0.1%) in four out of six opportunities, failing to detect the extremes in the *E. coli*/*S. saprophyticus* mixture. Reads matching phage present in the mixtures were classified and quantitated by Centrifuge separately from their host genomes. Because the phage relative abundance estimates were not included with their host, the bacteria present were underestimated so that the abundance estimates shown in Figure 1 do not add to 100%. The clearest example of phage matches affecting taxon-assignment is in the mixture composed of 99.9% *S. pyogenes* with an estimated relative *abundance of Streptococcus*-specific phage at 10.14%. Despite the effect of phage matches, the coefficient of determination (R^2^) for the three mixtures was 0.90 for *E. coli*/*S. flexneri*, 0.99 for *S. saprophyticus*/*S. pyogenes*, and 0.96 for *E. coli*/*S. saprophyticus*. Importantly, Centrifuge was able to discriminate between organisms as difficult to discriminate as *E. coli* and *S. flexneri*.

**Fig 1.**
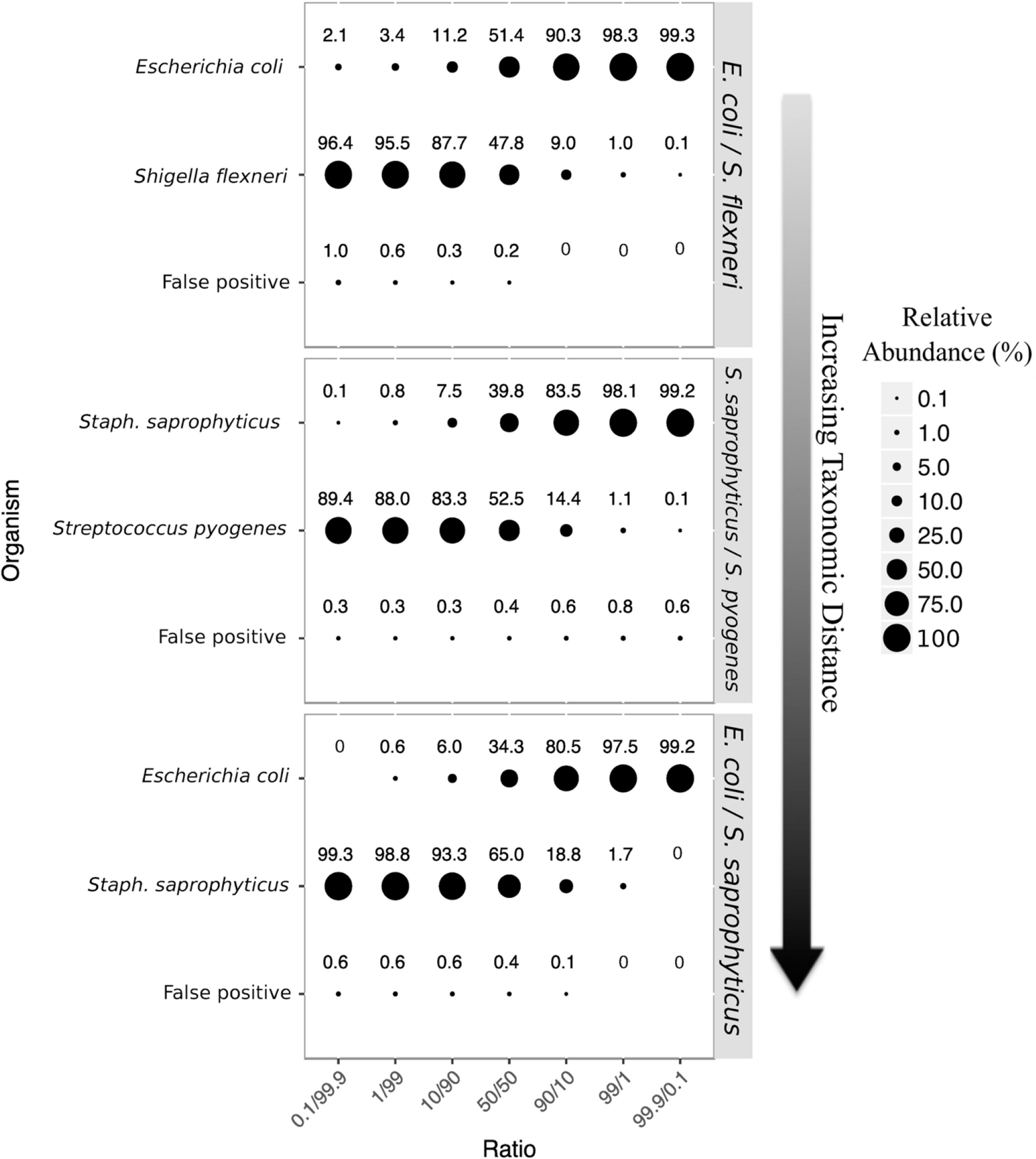
Linearity and threshold for detection of binary mixtures of bacteria using Centrifuge. The relative abundance of organisms calculated by Centrifuge is represented by circle size with actual values displayed above, values that are zero have no circle.

### Comparing the accuracy of Centrifuge and CLARK with a bacterial mock community

Given Centrifuge’s performance on the binary mixtures, it was next compared to a leading algorithm of another class, CLARK with a more complex mock community of 20 bacteria present in varying relative abundances. Both CLARK and Centrifuge identified the 20 known bacterial species in the mock community; however, CLARK reported five false positives (two *Shigella sp*., two *Staphylococcus sp*. and *Corynebacterium pseudotuberculosis*) that were not present in the mock community. In contrast to CLARK, Centrifuge did not produce any false positives. To compare the two algorithms (Centrifuge and CLARK), we graphed the relative abundance of 20 organisms in a mock community against their known abundance and calculated R2 values (Figure 2). Centrifuge and CLARK had nearly identical R^2^ values of 0.98 and 0.97 respectively. Overall, both tools tended to overestimate relative abundance values, especially the lowest abundances: most estimated abundances fell below the perfect fit represented by the dotted line in Figure 2. Importantly, both algorithms were able to identify the presence of all four organisms in the mock community with relative abundances of 0.01%.

**Fig 2.**
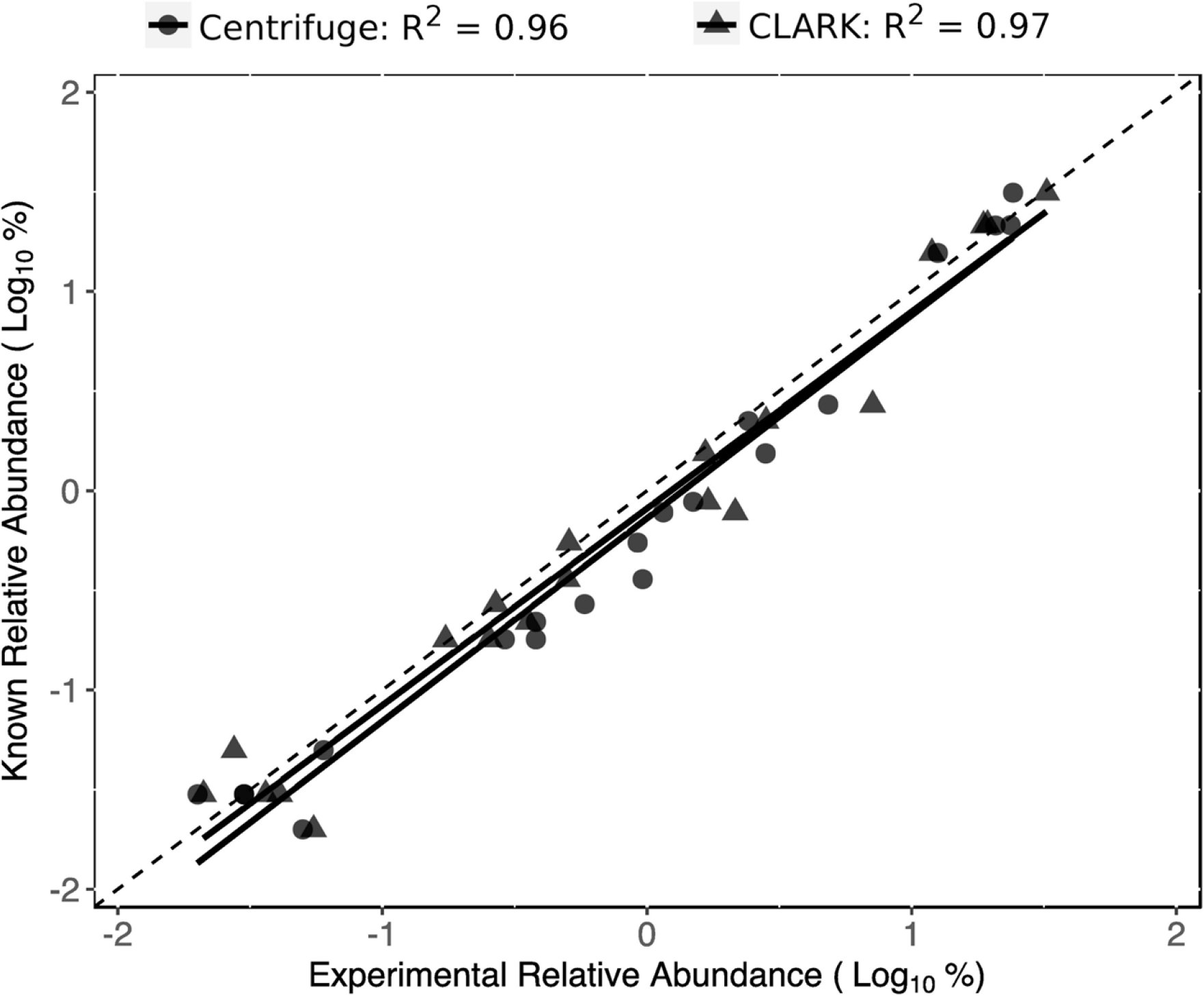
Centrifuge and CLARK relative abundance estimates versus expected for a mock community of 20 bacteria. Relative abundances estimated by CLARK and Centrifuge graphed against the expected values. The black dotted line represents perfect correlation with known relative abundances. The trendlines for CLARK and Centrifuge are shown in solid black lines.

### Centrifuge requires less computational resources than CLARK

While CLARK had nearly identical accuracy in relative abundance estimates as Centrifuge (despite five positive identifications), there was a striking difference between the two classification algorithms in the computation resources and time required to analyze the data. Relative to CLARK, Centrifuge required less than a tenth of the memory and a quarter of the runtime, while using half the number of central processing units (Table 1).

**Table 1.** Comparison of computational resources required by Centrifuge and CLARK to analyze the bacterial mock community dataset. CPU, central processing unit; GB, gigabyte; RAM, random access memory.

| Program | number of CPUs | RAM (GB) | Runtime (hr:min:sec) |
| --- | --- | --- | --- |
| Centrifuge | 12 | 23 | 0:07:40 |
| CLARK | 28 | 297 | 0:38:40 |

### Identification of pathogens in whole blood from febrile neutropenia patients

Pathogens were enriched using a simple sample preparation method from whole blood samples drawn from three patients with febrile neutropenia, and the resulting metagenomic DNA sequenced. Table 2 shows the starting number of raw reads and the percent passing through each step from quality control, to host-screening by alignment, and finally Centrifuge analysis. The reads classified by Centrifuge identified three likely pathogens: Pseudomonas fluorescens with a relative abundance of 50.7% in patient 1, Human parvovirus with a relative abundance of 99.8% in patient 2, and Torque teno virus in patient 3 with a relative abundance of 62.8% (Figure 3). Comparing the percentages shown in Table 2 with the relative abundances calculated by Centrifuge for these organisms showed how the small genome sizes of the two viruses gave their genomes more weight in the relative abundance estimates. For example, Torque Teno Virus had an abundance estimate of 72.8% though only 9.4% of the total post-quality control reads mapped to this organism.

**Fig 3.**
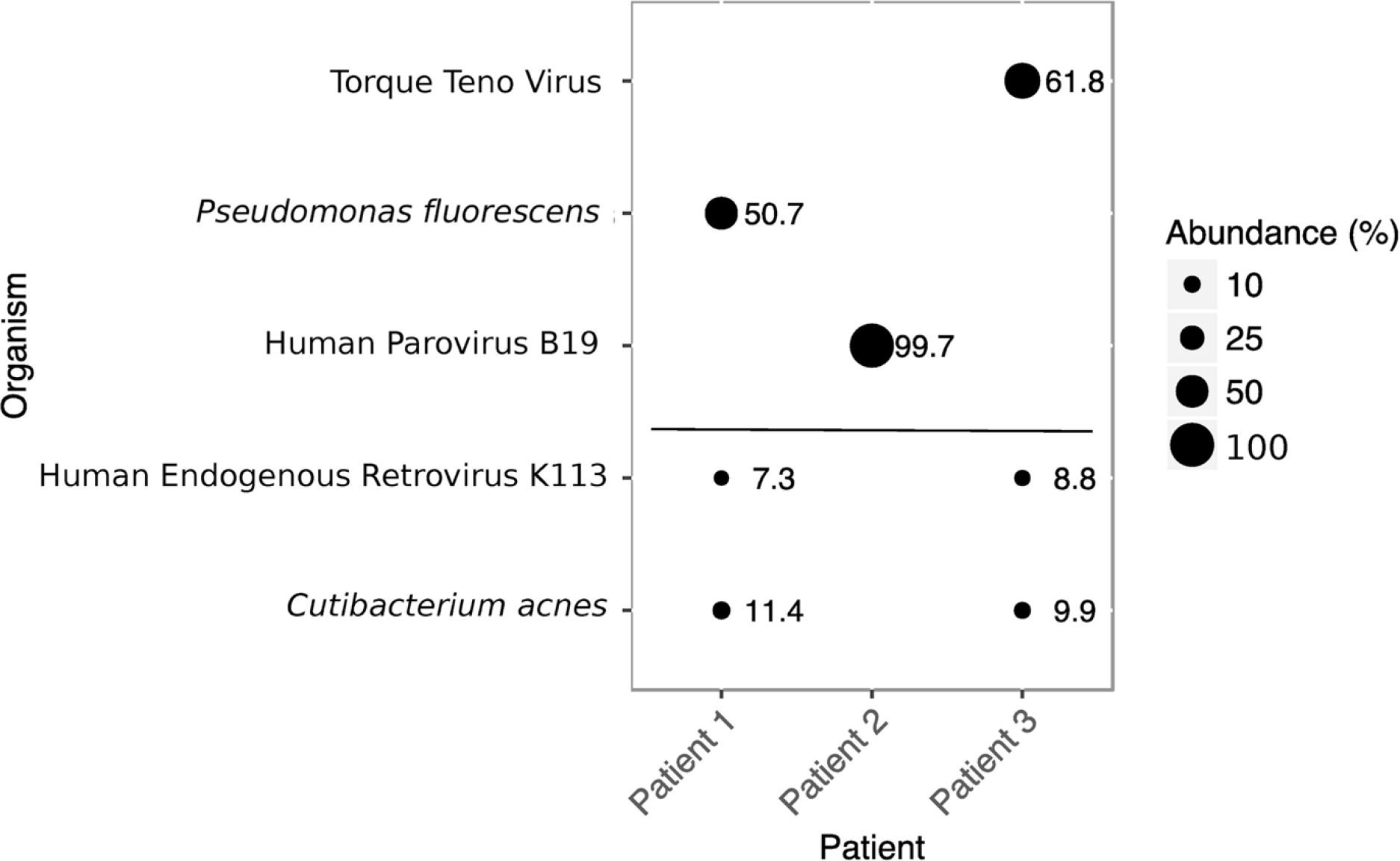
Identification and relative abundance of pathogens in febrile neutropenia samples. Circle size indicates the relative abundance of the respective organism, and actual abundance values are next to the circles. Organisms deemed endogenous or common contaminants are separated from the presumed pathogens by the horizontal line.

**Table 2.** Read counts following each step of the Centrifuge analysis of febrile neutropenia datasets. QC, quality control.

| Pt | Raw Reads<sup>a</sup> | Post QC<sup>b</sup> | Human (%)<sup>c</sup> | Centrifuge |  |  |
| --- | --- | --- | --- | --- | --- | --- |
|  |  |  |  | Unmapped (%)<sup>d</sup> | Classified (%)<sup>e</sup> | Unknown (%)<sup>f</sup> |
| 1 | 3,497,123 | 61.9 | 57.3 | 42.7 | 70.2 | 29.8 |
| 2 | 13,000,518 | 43.9 | 41.3 | 58.7 | 34.8 | 64.2 |
| 3 | 18,839,275 | 43.4 | 79.1 | 20.9 | 45.4 | 54.6 |

<sup>a</sup> Total reads generated from the sample.
<sup>b</sup> Percent of reads remaining after quality control.
<sup>c</sup> Percent of Post-QC reads that mapped to the human genome.
<sup>d</sup> Percent of Post-QC reads that did not map to the human genome.
<sup>e</sup> Percent of unmapped reads that were assigned a taxonomic classification by Centrifuge.
<sup>f</sup> Percentage of unmapped reads that were not assigned a taxonomic classification by Centrifuge.

Blood culture results for all three patients were negative, at the time of sample collection and in two subsequent blood cultures of each patient. Thus, the sequencing results were not compared to culture, the current gold standard. However, patient two did have a positive PCR test for human parvovirus in the month before and after the research sample was obtained, corroborating the results obtained with Centrifuge. Additional corroboration of the results comes from analysis of 12 samples obtained from two healthy volunteers over a six-week period in which none of the likely pathogens seen in the febrile neutropenia patients was observed (data not shown). While *Pseudomonas fluorescens* has been reported as a false positive in other studies, the fact that it did not appear in the healthy volunteer samples and is known to infect immunocompromised individuals (Wong et. al., 2011) suggests that it is not an artifact in patient 1 (. We also identified human endogenous retrovirus K113 and *Cutibacterium acnes* (also known as *Propionibacterium acnes*) in patients 1 and 3, however these organisms were deemed to be contaminants: the virus is endogenous, *C. acnes* is a common contaminant of blood samples (Mollerup et al., 2016; Parize et al., 2017; Park et al., 2011), and both were present in the normal samples collected over 6 weeks.

### Genome coverage of suspected pathogens in febrile neutropenic patients

Reads from the three febrile neutropenia samples were aligned to the respective reference genomes of the suspected pathogens to determine average depth of coverage (Figure 4). When patient 1 reads were aligned to the *Pseudomonas fluorescens* genome, the average coverage was 7.0. Patient 2 reads aligned to the Human Parvovirus B19 genome showed average coverage of 5,180. Finally, patient 3 reads aligned to the Torque Teno Virus (TTV) genome showed high coverage (~8,000) for a ~500 base pair region of the genome.

**Fig 4.**
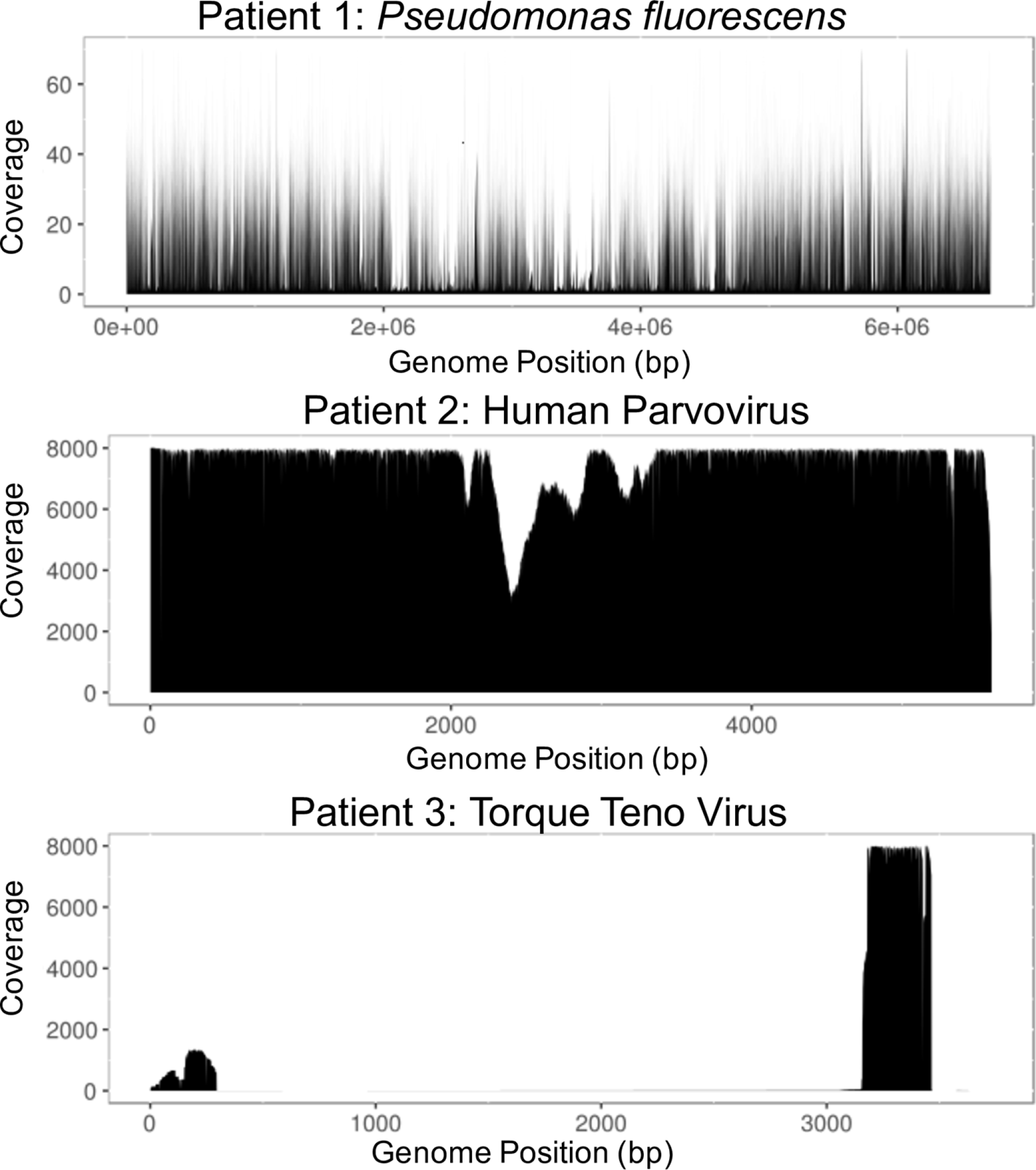
Genome coverage of suspected pathogens identified in febrile neutropenia patients by Centrifuge. Coverage by each base is graphed relative to the position in the three respective genomes of likely pathogens identified in three febrile neutropenia patients.

### Effect of quality controlling reads on computation time and Centrifuge’s accuracy

Sequencing reads are typically subjected to a series of quality control steps including trimming low-quality bases from reads, removing short reads, deduplication, and trimming ends with unbalanced nucleotide composition before downstream applications (e.g., variant calling, or sequence assembly). When quality control steps were performed before the Centrifuge analyses in Figures 2 and 3, they accounted for approximately half the compute time required to achieve results (data not shown). The fact that quality controls steps accounted for so much of the compute time, led to the question of what effect quality control had on the taxonomic classifications and relative abundance estimates made by Centrifuge. To answer this question, the mock bacterial community data was analyzed in Centrifuge with and without quality controlling the reads first. Results showed only one difference in taxonomic classification: a false positive (*Bacillus thuringiensis*) was identified with a relative abundance of 2.9% without quality control (Figure 5). Linear regression of the measured versus expected relative abundances showed that the R^2^ with quality control was 0.97 and without quality control was 0.97, further demonstrating how little effect there was on the Centrifuge results.

**Fig 5.**
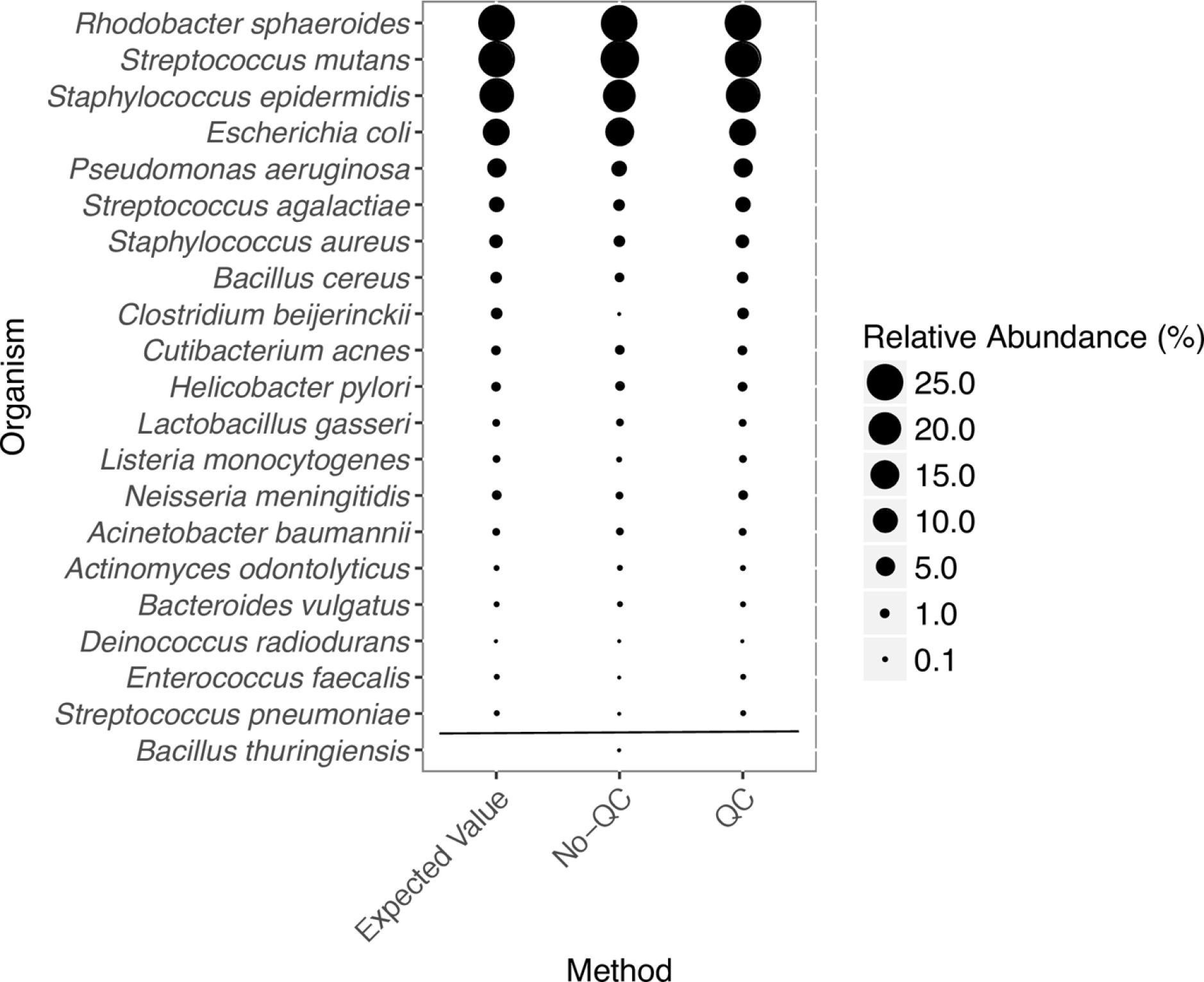
Bacterial mock community taxonomic identification and relative abundance by Centrifuge with and without quality control of the input sequence reads. Organisms are ranked by their relative abundance which is indicated by the size of the circle. The false positive (*Bacillus thuringiensis*) identified from reads without quality control (QC) is shown at the bottom.

### Host read removal by alignment versus in Centrifuge

Host DNA contamination can contribute to a significant proportion, or even the vast majority, of reads in metagenomic datasets, and is often removed by mapping reads to the host genome (Schmieder & Edwards, 2011). In performing taxonomic classification of reads, Centrifuge determines whether reads are of human origin (or other hosts), thus calling into question the necessity of aligning reads to the host genome and removing them, before analysis. Figure 6A shows the relative amount of reads that were classified as human, microbial, or unknown when the datasets were analyzed by Centrifuge without removing reads by alignment to the human genome before analysis. The relative proportion of host (human) reads in the data agreed well with the proportions found by alignment (see Table 2). While the proportion of host DNA was less than in prior studies, suggesting that the enrichment for pathogen DNA used in this study was successful, a significant proportion of the reads were still human.

**Fig 6.**
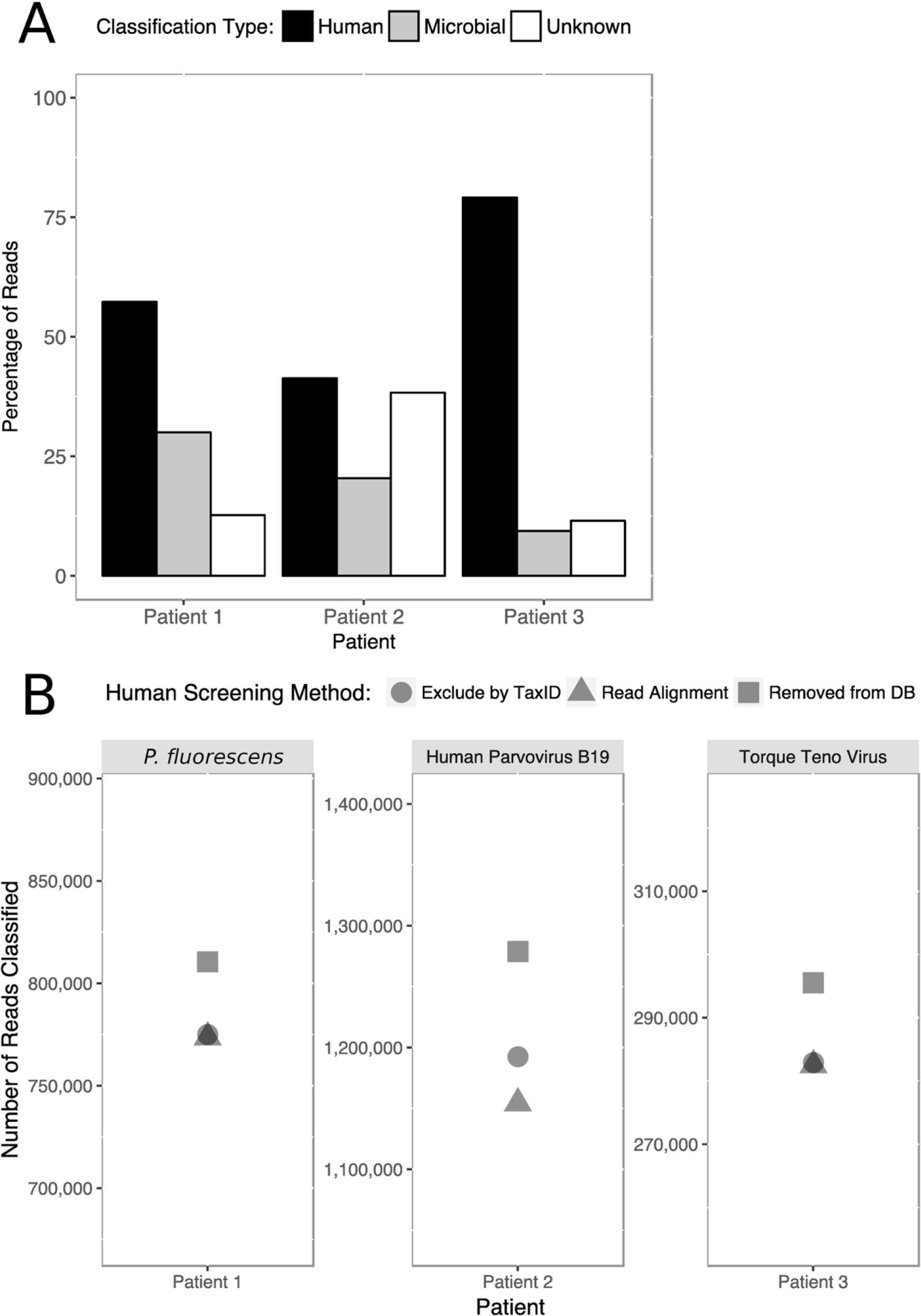
Effect of mapping reads from patient samples against the human genome. A) Percent of reads classified as human, microbial, or unknown for each febrile neutropenia patient by Centrifuge. B) Number of reads classified in each presumed pathogen following three strategies for host screening: removal of the human genome from the reference database (Removed from DB, squares), excluding the human TaxID in Centrifuge (Exclude by TaxID, circles), and aligning against the human genome before analysis (Read Alignment, triangles).

Having established that a significant proportion of the reads in the datasets were of host origin by both alignment and Centrifuge, we compared three approaches for removing host reads in the febrile neutropenia patient data. These methods include (1) alignment to the human genome and removal of aligned reads from the dataset, (2) removing the human sequence from the reference database, and (3) using the “exclude TaxID” function in Centrifuge to exclude reads from classification whose best match was to the human genome. Overall, exclusion of the human genome from the reference database resulted in the highest number of reads classified to the presumed pathogens; however, the differences between the methods were relatively minor (Figure 6B). Patient 3, with a presumed pathogen of Torque Teno virus, showed the least effect on the number of reads classified, with less than a 411 read difference (<1%) between the number of reads classified between the three methods. In contrast, patient 2, with a presumed pathogen of Human Parvovirus B19, had 124,544 fewer reads classified (9.7%) when reads removed by alignment relative to the removal of reads that match the human genome from the database. Finally, patient 1, with a presumed pathogen of *P. fluorescens*, showed 26,836 fewer reads classified (4.5%) when reads were removed by alignment relative to being present in the database.

## DISCUSSION

### Centrifuge accuracy of identification and quantitation with known samples

Immunocompromised patients, such as those with febrile neutropenia, are susceptible to infections. The current standard for identifying pathogens from clinical samples when infection is suspected can fail as much as ~80% of the time. Without diagnostic information, clinicians’ first response is empirical antibiotic therapy in the hope that the organism is bacterial and covered by the antibiotic(s) given. Metagenomic sequencing of clinical samples offers an approach that bypasses the issues of culture, however, mining the resulting metagenomic sequence can be slow and error-prone given the volume of reads, host read contamination, and lack of well-defined bioinformatics methods. The goal of our study was to assess Centrifuge, a leading tool for identification and quantitation of metagenomic data, using clinically relevant datasets to establish its accuracy in microbial/viral identification and abundance estimates with an eye toward reducing compute time.

The first dataset used to assess Centrifuge was a series of binary bacterial mixtures chosen for their phylogenetic distance and mixed so that each pair was combined across six logs of relative abundance. Centrifuge was able to discriminate the most closely related pair of bacteria, *E. coli* and *S. flexneri*, even when one of the organisms was present as 0.1% of the mixture. As the proportion of *E. coli* decreased, the relative abundance estimate diverged from expected, so that the *E. coli* estimate was 2.1% when *E. coli* was only 0.1% of the mixture. The same inaccuracy did not occur as the *S. flexneri* relative abundance decreased to 0.1%, suggesting Centrifuge misidentified a portion of the *S. flexneri* genome as *E. coli* but not the other way around. The difficulty classifying *S. flexneri* suggested by the fact that the false positive rate increased from 0% to 1%, the highest measured, as *S. flexneri* relative abundance increased. One likely cause for more relative matches to *E. coli* than *S. flexneri* is that *E. coli* strains and isolates represent the most substantial fraction of the Centrifuge reference database. False positive identification of *E. coli* using metagenomic methods has been previously observed. McIntyre et al. (2017) saw similar false positive identification of *E. coli* when using metagenomic classifiers on negative control sequences not belonging to any known organism(McIntyre et al., 2017). The researchers also speculated that the reason for the false positives is the overrepresentation of *E. coli* sequences in their reference dataset. Although Centrifuge uses a modified FM-index to condense closely related genomes, the total file size of basepairs maintained (unique + shared based on ≥ 99% identity) exceeds the relative file size of all other species (Kim et al., 2016) giving it a higher probability for matches. This result suggests that Centrifuge dampens the effect of multiple strains and isolate genomes using the modified FM-index, but the effect is still present for highly abundant strains.

Centrifuge appears to be capable of detecting organisms even when they are present in minor abundance, regardless of the phylogenetic distances between them. Overall, Centrifuge read abundances closely match the expected relative abundance of bacterial mixtures for closely and distantly related species. Interestingly, phylogenetic distance did not predict the accuracy of relative abundance estimates. A reasonable assumption would be that as phylogenetic distance increases, the number of discriminatory k-mers increase to allow for better read classification by Centrifuge. Instead, we observed high classification accuracy for the most closely related pair (*E. coli*/*S. flexneri)* from the same family. Less accuracy for the next pair (*S. pyogenes*/*S. saprophyticus*) where both organisms were gram-positive and from the same phylogenetic class. The highest accuracy for the most distant pair (*E. coli*/*S. saprophyticus*) where one organism was gram-negative and the other gram-positive and only shared phylogenetic kingdom. Interestingly, *S. pyogenes* is closely related to many *Streptococcus* genomes which may have limited the number of distinguishing k-mers to classify reads at the species rather than genus level (data not shown).

We compared Centrifuge’s performance against another leading k-mer based taxonomic classifier, CLARK, in analyzing sequence data from a more complex community of 20 bacteria. The mock community was also mixed in varying relative abundances as with the binary mixtures, albeit, in a different range (~0.01-35%). Abundance calculations between the two algorithms were nearly identical across the relative abundance range; however, the processing time and computational resources for CLARK were greater (Table 1). Also, CLARK had a propensity for false positives, whereas Centrifuge did not. On the other hand, Centrifuge’s results had to be processed to account for the strain and phage-specific data generated. Such processing would be a necessary part of adoption in a clinical setting, but Centrifuge’s lack of false positives and speed suggests it may be a good starting point for such a tool.

### Centrifuge identification and relative abundance estimates

Centrifuge is unique from other taxonomic classifiers in that it provides Expectation – Maximization (EM) calculation to determine relative abundance, rather than just read proportional classification. The EM calculation proves useful in determining relative abundance between organisms in samples with varying genome sizes. We demonstrated the benefit of calculating abundance using Centrifuge’s EM algorithm in the analysis of the febrile neutropenia blood samples from patients 2 and 3 where viral matches were significantly underrepresented when using read proportional classifications.

One drawback for clinical pathogen identification is that Centrifuge separates strain-level counts, splitting reads among closely related strains which required manually summing strain level abundances for reporting. Future iterations of Centrifuge could address this issue re-analyzing the data with a reduced reference set of genomes based on the first round of analysis or a reduced reference database. Lastly, current reference databases do not account for all of the extant microbial/viral diversity that may be present in patients. However, this issue is being addressed over time with the exponential growth in the number of microbial draft genomes available (Land et al., 2015).

### Genome coverage of presumptive pathogens identified in patient samples

We examined genome coverage statistics with the assumption that the genomes of the pathogens identified as the presumed cause of fever in the patients would be represented by consistent coverage, whereas uneven coverage could indicate insufficient evidence of organism presence. Parize et al. took a similar approach in which even distribution of contigs was used as part of the criteria to decide if a sample was deemed positive (Parize et al., 2017). Interestingly, the Torque Teno virus sequence found in patient 3 was observed to have high coverage of only a ~500 base pair untranslated region of the genome. This highly conserved region has been suggested to be critical for viral replication that may indicate an early replication event or the presence of subviral particles, a characteristic that has previously observed in Torque Teno virus (de Villiers, Borkosky, Kimmel, Gunst, & Fei, 2011). The evidence for sub-viral particles provided by the coverage analysis is the first from an *in vivo* sample. Lastly, Torque Teno virus was identified in a cancer patient undergoing bone marrow ablation in preparation for a hematopoietic stem cell transplant as part of their cancer treatment. This finding highlights the possible value of the metagenomic sequencing approach as Torque Teno virus has been investigated as a predictive marker for post-transplant complications (Wohlfarth et al., 2018).

### Quality control of reads before Centrifuge analysis

Although quality control of raw reads is imperative for variant calling and genome assembly and can speed up downstream taxonomic and functional analyses by reducing the total number of reads analyzed, it takes considerable computing time and resources. In this study, we observed limited benefits of quality control regarding accurately identifying and quantifying the abundance of the bacteria in the mock community. However, we did see an elimination of a single false positive organism estimated at 2.3% relative abundance with quality control. Quality controlling reads from the febrile neutropenia data revealed a bias toward removing viral reads (Supplemental Table 1). Users of Centrifuge may want to weigh the limited benefits of quality controlling their data before analysis in Centrifuge versus the bias toward the removal of viral reads and time required.

### Host screening with Centrifuge

Despite the substantial enrichment for microbial/viral DNA that we achieved in this study (20-58% non-human reads, Table 2) as compared to prior studies (1% of reads) (Naccache et al., 2014; Parize et al., 2017), a large proportion of reads were still identified as human. Screening host reads by alignment to the genome before analysis by Centrifuge appears to be unnecessary given Centrifuge’s ability to classify reads to the host organism during analysis. For example, in patient 2 we were able to identify Human Parvovirus B19 when we used the “exclude TaxID” function for host screening. Because parvovirus virus integrated into the ancestral human genome during evolution (Liu et al., 2011), many Human Parvovirus B19 reads identified aligned to the human genome and were removed before analysis by Centrifuge. This method caused the largest reduction in the number of reads classified as Human Parvovirus B19 relative to the exclude TaxID method (Figure 6B).

In contrast, when the human genome was removed from the Centrifuge database, reads from the human genome derived from the ancestrally integrated parvovirus would have been misclassified as Human Parvovirus B19, with the effect that it could inflate the relative abundance estimate. The “exclude TaxID” method appears to offer a balance between the other two methods: it allows both endogenous host reads and actual organism reads to be appropriately classified while saving the time and computational cost of aligning reads to a host organism before analysis. Given that reference genomes can contain sequences of mixed origin due to horizontal gene transfer, endogenous and integrated microbes/viruses, and prophage in bacterial genomes, classifying reads to all available reference data and then utilizing exclude TaxID appears to be the best compromise of speed and specificity for eliminating host reads from results.

## Conclusion

In summary, our analyses suggest that Centrifuge, open-source software for fast taxonomic classification, provides accurate quantification of clinically relevant organisms/viruses in metagenomes using minimal compute time and resources. Centrifuge’s ability to quickly assign taxonomy to reads, accurately represent the abundance of organisms such as viruses, and sidestep read quality control and host-screening make it a good candidate for classifying reads of clinically relevant organisms. To this end, we have made Centrifuge and the bubble plot software used in the study available as Apps in iMicrobe (http://imicrobe.us) for streamlined taxonomic analysis by the public.

## Materials and Methods

These methods have been deposited into protocols.io under DOI: dx.doi.org/10.17504/protocols.io.wjdfci6

### Ethics Statement

The Institutional Review Board at the University of Arizona (project #1505826794) approved the human subjects research. Informed consent was obtained from febrile neutropenia patients. Whole blood was collected from patients that developed febrile neutropenia during their treatment at the University of Arizona Cancer Center. Data obtained from the first three patients collected as part of a more extensive study were used here. All three patients were being treated for leukemia or lymphoma at the time of their febrile neutropenia diagnosis.

### Binary mixtures of bacteria

The binary mixtures were described previously (Watts et al., 2017). Briefly, four species of bacteria were used to create three binary mixtures representing: (1) difficult to discriminate species with divergent clinical impact (*Escherichia coli* versus *Shigella flexneri*); (2) Gram-positive species (*Staphylococcus saprophyticus* versus *Streptococcus pyogenes*); and (3) Gram-positive versus Gram-negative species (*E. coli* versus *S. saprophyticus*). DNA from the bacteria were purchased from the American Type Culture Collection (Manassas, Va, USA) and mixed in pairs so that each species represented 99.9, 99, 90, 50, 10, 1, and 0.1% of the total sample. Samples were sequenced as described below, and the sequence data deposited to the NCBI Sequence Read Archive under accessions: SRX3154186-SRX3154219 in project accession PRJNA401033.

### Staggered mock bacterial community

The mock bacterial community (BEI Resources, Manassas, VA, USA, National Institute Allergy and Infectious Diseases, National Institutes of Health, as part of the Human Microbiome Project: Genomic DNA from Microbial Mock Community B (Staggered, High Concentration), v5.2H, for metagenomic shotgun sequencing, HM-277D) consisted of 20 bacterial species created as part of the Human Microbiome Project with specific staggered 16S rRNA gene abundances for each species. Using the 16S rRNA gene copy values, along with the known 16S rRNA gene copy number in each species’ genome, we calculated the number of genomes present for each species to provide an expected value for comparison to the relative abundances calculated by Centrifuge and CLARK from sequencing data. The mock community was sequenced as described below, and sequence data deposited to the NCBI Sequence Read Archive under accession: SRP115095 in project accession PRJNA397434.

### Febrile neutropenia patient blood samples

Approximately five milliliters of whole blood were collected (K_2_EDTA BD Vacutainer tubes, catalog #367863 BD Biosciences, San Jose, CA, USA) when blood cultures were ordered for each patient and transferred for processing within 2 hours of collection. Blood samples were diluted with an equal volume of sterile phosphate buffered saline, layered on Ficoll-Paque (GE HealthCare Life Sciences, Pittsburgh, PA, USA) and centrifuged for 20 minutes at 400 x g. Plasma was carefully drawn off, sacrificing some yield to prevent drawing up monocytes, and centrifuged three more times at 50, 100, and 150 x g for 5 minutes to further remove human cells. The plasma was passed through a five-micron filter and finally centrifuged at 4000 x g. DNA was isolated from any material sedimented during the final centrifugation with a UCP Pure Pathogen kit (Qiagen Inc., Germantown, MD, USA). Isolated DNA was quantitated on a NanoDrop ND-1000 spectrophotometer at 260 nanometers (Thermo Fisher Technologies Inc., Santa Clara, CA, USA), diluted to one nanogram/microliter, and ten nanograms used to prepare sequencing libraries as described below. Sequence data for the three patient samples were deposited to the NCBI Sequence Read Archive in project accession PRJNA521396.

### DNA library preparation and sequencing

DNA libraries were prepared and sequenced for all samples utilizing Ion Torrent reagents and the Ion Torrent Proton sequencer (Thermo Fisher Technologies Inc., Santa Clara, CA, USA). Ten nanograms of DNA was input to the Ion Xpress Plus Fragment Library Kit (manual #MAN0009847, revC). DNA was sheared using the Ion Shear enzymatic reaction for 12 min, and Ion Xpress barcode adapters were ligated following end repair. Resulting libraries were amplified using the manufacturer supplied library amplification primers and recommended conditions. Amplified libraries were size selected to approximately 200 base pairs using E-gel SizeSelect Agarose cassettes (Invitrogen, Carlsbad, CA, USA) as outlined in the Ion Xpress manual and quantitated with the Ion Universal Library quantitation kit. Equimolar amounts of the library were templated with an Ion PI Template OT2 200 kit V3. The resulting templated beads were enriched with the Ion OneTouch ES system and quantitated with the Qubit Ion Sphere Quality Control kit on a Qubit 3.0 fluorimeter (Qubit, NY, NY, USA). Enriched templated beads were loaded onto an Ion PI V2 chip and sequenced according to the manufacturer’s protocol using the Ion PI Sequencing 200 kit V3. Data were processed with Ion Torrent Server software v4.4.3 to produce data files in BAM format.

### Read processing and quality control

Sequences were converted to FASTQ format from raw BAM files with bedtools’ bamtofastq (Quinlan & Hall, 2010)2.17.0, (Quinlan & Hall, 2010). FastQC (“Babraham Bioinformatics - FastQC A Quality Control tool for High Throughput Sequence Data,” n.d.) v0.11.5, (“Babraham Bioinformatics - FastQC A Quality Control tool for High Throughput Sequence Data,” n.d.) was used to generate sequence quality reports. FastX toolkit (Gordon & Hannon, 2010)v.0.0.14, (Gordon & Hannon, 2010) was used to perform quality control measures on FASTQ data including quality filtering, trimming, setting a minimum read length, and removal of duplicate reads. Files were converted to FASTA with FastX. Data files before and after QC were used as input to Centrifuge when testing the effect of quality control; otherwise, all files were quality controlled before analysis.

### Removing host contamination by aligning to the human genome

To remove host (human) reads, FASTQ read files were mapped to HG38 (Genome reference consortium human genome build38) using Bowtie2 (Langmead & Salzberg, 2012) using the --very-sensitive option. Human reads were removed by alignment from patient data before the analysis in Centrifuge except when testing the effect of host screening by other methods.

### Centrifuge and CLARK read classification

CLARK v1.1.3 (Ounit et al., 2015a) was used to classify reads to known taxa using the default CLARK database and parameters. Centrifuge v1.0.3-beta (Kim et al., 2016) was used to classify reads to known taxa with a custom database generated from 23,276 complete archaeal, bacterial, and viral genomes downloaded from Refseq in July 2017 using the centrifuge-download and centrifuge-build scripts respectively. The custom database is available at https://github.com/hurwitzlab/NeutropenicFever.

### Binary mixture Centrifuge results filtering

Centrifuge abundance report results were filtered to only include organisms at the species or strain-level with a minimum of 0.1% of total reads classified and at least 0.05% abundance as calculated by Centrifuge. These settings were chosen based on the known abundances used in the mixtures. False positive was calculated by summing the relative abundances of any organism identified by Centrifuge that was not added to the mixture. Centrifuge reports read-matches to phage separately from their host species; however, no phage or prophage passed the above filters, so there was no effect on the relative abundance calculations for the binary mixtures. The coefficient of determination (R^2^) was calculated based on the log of both relative abundance estimates at each known dilution.

### Mock community Centrifuge results filtering

Centrifuge abundance report results were filtered to only include organisms at the species or strain-level with a minimum of at least 0.005% abundance as calculated by Centrifuge and no minimum number of reads. These settings were chosen based on the known abundances calculated for the mock community which was lower than the bacterial mixtures (0.01%). In the case of the mock community, two species-specific phages were identified that passed the filters (*Pseudomonas* phage with relative abundance 1.5%, and *Staphylococcus* phage with relative abundance 0.8%). The matches to these phages were included when calculating relative abundances for the 20 organisms, but not included in the figure. The coefficient of determination (R^2^) was calculated based on the log of the relative abundance estimates for all 20 species.

### Febrile Neutropenia Centrifuge results filtering

Centrifuge abundance report results were filtered to only include organisms at the species or strain-level with a minimum of 1% of total reads classified and at least 5% abundance as calculated by Centrifuge. Similarly to the bacterial mixtures, no phage or prophage passed the filters above, so there was no effect on relative abundance calculations.

### Genome coverage of suspected pathogens from febrile neutropenia patient samples

To determine genome coverage, we used Bowtie2 (Langmead & Salzberg, 2012) to map FASTQ reads (with option --very-sensitive) to reference genomes for the organisms identified by Centrifuge (*Pseudomonas fluorescens* accession: NC_012660.1, Human Parvovirus B19 accession: NC_000883.2, Torque Teno Virus accession: NC_015783.1). Resulting BAM files were then analyzed utilizing Samtools’ (v1.3.1, (Li et al., 2009) depth tool to generate coverage values and visualized in R v3.1.1 (R scripts are available here: https://github.com/hurwitzlab/NeutropenicFever).

### Software availability

To improve access to Centrifuge and the bubble chart visualizations used in this manuscript, both tools have been made available on iMicrobe (https://www.imicrobe.us). As a starting point, researchers may run <u>centrifuge-0.0.6u1</u> followed by <u>centrifuge-bubble-0.0.5u1</u> to reproduce the bacterial mixing results in the manuscript using the sample data provided. Source code for running centrifuge on a high-performance compute cluster is available in Github at https://github.com/hurwitzlab/Centrifuge_HPC and analyses, scripts and visualizations are also archived at https://github.com/hurwitzlab/NeutropenicFever.

## ACKNOWLEDGMENTS

We thank Drs. Jana U’Ren and Alise Ponsero for feedback on the manuscript. We thank Dr. Marvin Slepian for constructive discussions. We thank the staff of the University of Arizona Information Technology Services for access and system support for the UA high-performance compute cluster. We thank The University of Arizona BIO5 Statistical Consulting Services for advice on calculating the coefficients of determination.

## SUPPORTING INFORMATION

**S1. Quality Control Reduction of Reads**

